# Confidence in action: differences between perceived accuracy of decision and motor response

**DOI:** 10.1101/2020.06.03.132068

**Authors:** Marta Siedlecka, Marcin Koculak, Borysław Paulewicz

## Abstract

Each of our decisions is associated with a degree of confidence. This confidence can change once we have acted as we might start doubting our choice or even become convinced that we made a mistake. In this study, we explore the relations between action and our confidence that our decision was correct or erroneous. Fifty-six volunteers took part in a perceptual decision task in which their decisions could either lead to action or not. At the end of each trial, participants rated their confidence that their decision was correct, or they reported that they had made an error. The main results showed that when given after a response, confidence ratings were higher and more strongly related to decision accuracy, and post-response reports of errors more often indicated actual errors. The results support the view that error awareness and confidence might be partially based on post-action processing.

## Introduction

Each of our decisions is associated with a degree of confidence, but this confidence can change once we have acted as we might start doubting our choice or even become convinced that we made a mistake. In the study reported in this paper, we explore the relations between action and our confidence that our decision was correct or incorrect.

Confidence in decisions and the ability to recognize our errors are manifestations of metacognition, i.e. processes that monitor and regulate other cognitive processes. Interestingly, studies on confidence and error awareness have mostly been run in parallel. In a typical study on confidence, participants make a series of forced choices about external stimuli or their memory content; subsequently, they rate their confidence. In such tasks, confidence is typically measured with a scale ranging from uncertainty to certainty in a given choice. Participants rate their confidence in having responded correctly, and we have no knowledge about what rating they choose when they know they made a mistake. In studies on error awareness, forced-choice tasks are also used; they are usually easier than those used in confidence studies, but they elicit more errors due to time pressure. Importantly, in studies on error awareness participants do not typically report graded confidence; instead, they are asked to signal when they think they made a mistake or to classify each response as correct or incorrect (Rabbit, 1968; Steinhauser & Yeung, 2012; Wessel, Danielmeier, & Ullsperger, 2011). To our knowledge, only a few studies have measured both confidence in being right and in being wrong, i.e. by using a scale ranging from “certainly wrong” to “certainly correct” (Boldt & Yeung, 2015; Charles & Yeung, 2019; Scheffers & Coles, 2000). Confidence measured in this way has been shown to covary with neural activity associated with error commitment, thus leading some authors to suggest that error reports and confidence judgments result from the same metacognitive processes (Boldt & Yeung, 2015; Charles & Yeung, 2019).

What is the relation between action and metacognitive judgments? Studies have shown that confidence better predicts response accuracy when it is measured after a decision-related response (Pereira, Faivre, Iturrate, Wirthlin, Serafini, Martin, Desvachez, Blanke, Van De Ville, & Millán, 2018; Siedlecka, Skóra, Fijałkowska, Paulewicz, Timmermans, & Wierzchoń, 2019; Wokke, Achoui, & Cleeremans, 2019). Interpreted in the context of the observed relations between confidence level and action characteristics (Fleming, Maniscalco, Ko, Amendi, Ro, & Lau, 2015; Susser & Mulligan, 2015), these results led to the conclusion that response-related signals inform confidence judgments. However, in these studies motor responses were indistinguishable from decisions, making it difficult to determine whether it is the action or the choice itself that improves metacognitive assessment (Kvam, Pleskac, Yu, & Busemeyer, 2015). On the other hand, when the type of the action that precedes a confidence judgment is manipulated experimentally, then certain types of responses (e.g. compatible with the decision-related stimulus characteristics) are shown to increase confidence (Filevich, Koß, & Faivre, 2019; Siedlecka, Paulewicz, & Koculak, 2020) without affecting the relation between confidence ratings and decision accuracy (Filevich et al., 2019).

While confidence is usually measured as an assessment of decision accuracy (inferred from motor responses), errors are almost exclusively studied in the context of action control and performance monitoring. These studies are focused on how action selection and execution is evaluated and regulated; therefore, motor errors are of most interest. These errors are generally divided into two types: slips, which are unintended actions (Norman, 1981); premature responses, which are responses that are given before stimulus processing has completed (Rabbitt, 2002). Such errors could be detected by the monitoring system by registering an action plan that is in competition with an already launched response (Rabbitt & Vyas, 1981) or by detecting conflict between several activated responses (Yeung, Botvinick, & Cohen, 2004). However, such errors are often unnoticed and unreported as the processes of error detection and correction can be relatively quick and automatic. Error awareness, on the contrary, is thought to result from a post-action, time-consuming accumulation of information from numerous sources, including sensory and proprioceptive feedback from the erroneous action, and interoception of the autonomic responses that accompany an error (Ullsperger, Harsay, Wessel, & Ridderinkhof, 2010; Wessel, 2012). However, it is not clear whether and how people can make accurate judgments about non-motor-type errors: for example, when an incorrect decision is not associated with any action.

In the presented study we compared metacognitive judgments between trials in which decisions were followed by action and in which they were not. We combined paradigms used for measuring confidence and error awareness in a task that allowed separation of decision and motor responses. One difficulty in testing the influence of response on metacognitive judgments is that manipulating the presence of motor response does not allow the accuracy of decisions that are not expressed with a behavioural response to be precisely assessed. In this task, participants were asked to act or not act, depending on their decision about the stimuli. This allowed identification of their covert choices, even when they did not overtly respond. Assuming that both types of metacognitive judgments could be affected by action-related information, we hypothesized that confidence and error reports would be more accurate following motor responses. We also expected higher confidence in decisions that were followed by actions.

## Methods

### Participants

Fifty-four healthy student volunteers (6 males), aged 19–32 (M = 20.72, SD = 2.69) took part in the experiment in return for credit points in a Cognitive Psychology course. All participants had normal or corrected-to-normal vision and gave written consent to participate in the study. The Ethical Committee of the Institute of Psychology, Jagiellonian University approved the experimental protocol. Student participants were chosen because this is a group that is relatively easily available to the researchers at our University and at the same time these participants are used to solving computer tasks. Because of the task novelty, we were not able to estimate sample size, so we decided to test a reasonably large group during available laboratory time (around 50 participants).

## Materials

We used a perceptual decision task in which participants are asked to decide which of the two presented fields, left or right, contains more dots; then they rate their confidence in this response (Boldt & Yeung, 2015, Siedlecka et al., 2020). However, the task was modified such that it required the spacebar to be pressed when there were more dots on one side, and participants were instructed not to respond when there were more dots on the other side. In one block, participants used their left thumb to indicate a “left” decision and did nothing for “right” decisions, while in the other block they used their right thumb to indicate a “right” decision and did nothing for “left” decisions. This design allowed us to divide trials into two groups (later called conditions): those in which participants responded (Response condition) and those in which participants did not respond (No response condition). At the end of each trial, participants were asked to rate their confidence in their choice or report an error. The outline of the experimental trial is presented in Figure 1.

**Figure 1.**
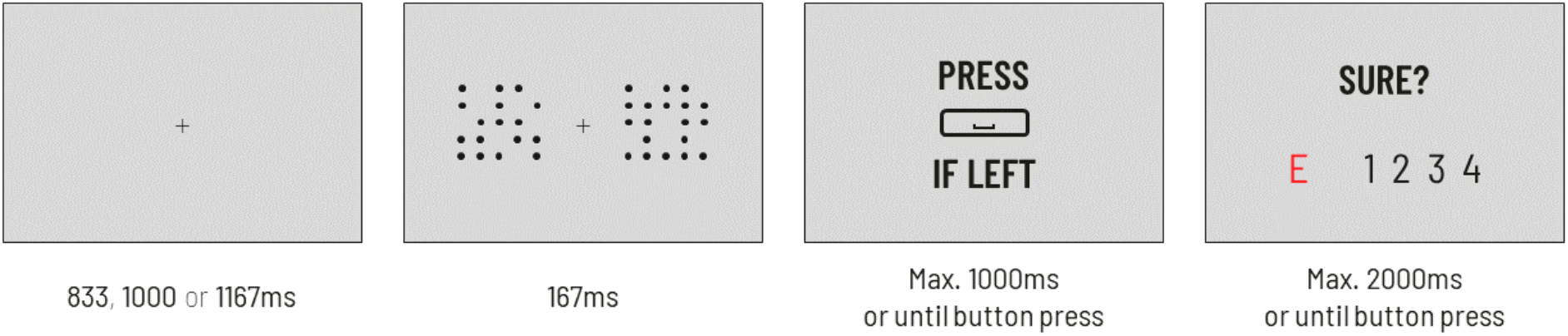
Outline of the experimental trial. First, a fixation cross was displayed, followed by a brief presentation of the grid stimuli. Immediately after that, participants were asked to decide if there were more dots on one side (here: left) and to report this decision by either pressing the spacebar (here: if they decided “left”) or not responding (here: if they decided “right”). Finally, participants rated their confidence in the correctness of their decision on a 4-point scale; alternatively, they could report that they had made an error.

The experiment was run on a PC using PsychoPy software (Pierce, Gray, Simpson, MacAskill, Höchenberger, Sogo, Kastman, & Lindeløv, 2019) with an LCD monitor (1920 x 1080 pixels resolution, 60-Hz refresh rate). Experimental stimuli were composed of two 10 x 10 grids filled with dots of equal size (160 by 160 px). Grids were placed at an equal distance of 120 pixels from the centre of the screen in the horizontal plane. In each grid, a predefined number of dots was placed randomly in each trial. Based on our previous pilot studies with the dot task (with responses for both decisions), the proportions of dots chosen for the experimental session were estimated to lead to an overall accuracy of 70%.

Participants responded by pressing buttons on a standard PC keyboard. The space bar was used for the response in the decision task. To measure participants’ confidence, we used a four-point scale: 1 – “I am guessing”, 2 – “I am not confident”, 3 – “I am quite confident” and 4 – “I am very confident”. Participants used their right hands and the keys U, I, O, P, respectively (labelled 1, 2, 3, 4 in Figure 1). Additionally, participants could report that they had made an error by pressing the H key (labelled E in Figure 1).

### Procedure

Participants were tested individually in a soundproof cabin with fixed lightning conditions. After receiving the general instructions, participants were told to place their fingers on the response keys so they would not look at the keyboard during the task. Participants were instructed to respond as accurately as they could. They were also told that the number of trials with correct “left” and “right” answers was equal in each block. Participants were explicitly asked to avoid refraining from responding when they were not confident about the correct decision, and they were told that after every 30 trials they would receive feedback about their accuracy and the percentage of trials in which they responded. Lastly, the experimenter asked participants to try to differentiate their confidence levels between trials even if they felt that the task was generally difficult.

The experiment consisted of three parts: two training sessions and the main task. A trial started with a fixation cross presented centrally on the screen for around one second; this was followed by a brief presentation of the grid stimuli with dots. After the stimuli disappeared, participants saw a screen prompting them to make a decision about the stimuli and to respond or not, according to whether the current block required the “left” or “right” decision to be reported. Finally, a screen with a confidence scale appeared on which participants rated their confidence in their decision or could signal that their decision was incorrect.

The first training session had a relatively low difficulty level as it was designed to accustom participants to the experimental procedure. This was achieved through presenting the grids with dots for 334 ms with a fixed dot ratio of 70:30 and 60:40. Participants were allowed 2.5 seconds for stimuli-related decisions, and 3.5 seconds were allowed for the confidence rating and error report. Subsequently, participants went through a second training session which looked like the main task except for the ratio of the dots (55:45). We introduced variable fixation cross presentation time (833, 1000 or 1167 milliseconds) to prevent perceptual entrainment and minimize the effects of expectations. In the main task, the ratio of the dots between the grids was changed and three levels of decision difficulty were introduced: 55:45, 52:48 and 51:49. The time available for the decision was 1 second, while 2 seconds were allowed for the confidence rating and error report.

There were two blocks in the main experimental task: responding when the decision was “left” and responding when the decision was “right”. The order of blocks was counterbalanced. The two blocks were bundled into 13 series of 30 trials (summing up to 390 trials per block and 780 trials in the whole experiment). After each series, the participant was provided with a feedback screen (accuracy and proportion of left/right responses) which lasted at least 10 seconds, but they could extend this break if needed.

During this procedure, EEG and EMG recordings were also acquired but these were unrelated to the hypotheses tested in the current study and are not presented here.

## Results

Statistical analyses were run in the R statistical environment (R Core Team, 2019) using linear or logistic mixed regression models. The models were fitted using the lme4 and lmerTest packages (Bates, Maechler, Bolker, & Walker, 2015; Kuznetsova, Brockhoff, & Christensen, 2017). Series of models with all possible random effects and their interactions were fitted to estimate confidence level and response times (a linear mixed model) and accuracy (a logistic mixed model).

### Data set preparation

The trials in which participants did not rate their confidence were omitted from analysis. We also excluded the data of one participant that differed in scale-use variance from the rest of the group (based on visual inspection of the sorted confidence-rating variances scores plot). The data of 53 participants (38,733 observations) were included in the final dataset. Three participants did not succeed in adapting to the new instruction (i.e. responding in the case of a “left” decision in the current block after they were supposed to respond in the case of a “right” decision in the previous block) and their performance in the second block was at or below chance level. The data of these three participants from the second block were not included in the analyses.

### Performance in the decision task

First, we compared the accuracy between trials that required “left” and “right” decisions. A mixed logistic regression model with random effect of condition and participant-specific intercept did not detect statistically significant differences (*z* = −0.51, *p* = .61). Fitting a mixed logistic regression model with random intercept revealed that participants more often chose not to respond than to respond (*z* = −2.92, *p* < .05). Moreover, when the fixed and random effect of Response was included, the results indicated that participants’ accuracy was higher when they responded compared to trials in which they did not respond (*z* = 2.99, *p* < .01). The general statistics related to task performance are presented in Table 1.

**Table 1.** Decision accuracy, number of trials, number of error reports, and error rate when reporting an error in the Response and No response conditions.

|  | Decision accuracy | Number of trials | Frequency of error report | Error rate when reporting error |
| --- | --- | --- | --- | --- |
| **Response** | 72% | 18,506 (48%) | 535 (2.9%) | 84% |
| **No response** | 70% | 20,227 (52%) | 501 (2.5%) | 56% |

Signal detection analysis using mixed logistic regression with the probit function revealed that Response trials were associated with higher discrimination sensitivity (*z* = 2.51, p < .05), but we did not detect significant differences in response bias (*z* = −0.55, *p* = .58).

### Error reports

A mixed logistic model with fixed and random effect of condition and participant-specific intercept did not reveal a significant difference in the number of reports between the conditions (*z* = 1.85, *p* = .06). However error reports were more accurate when they followed motor response (Table 1): a mixed logistic model with fixed and random effect of condition and participant-specific intercept fitted to data from trials in which an error was reported revealed that when an error was reported in Response trials, the actual error rate was significantly lower in No-response trials (*z* = −7.48, *p* < .001, Figure 2).

**Figure 2.**
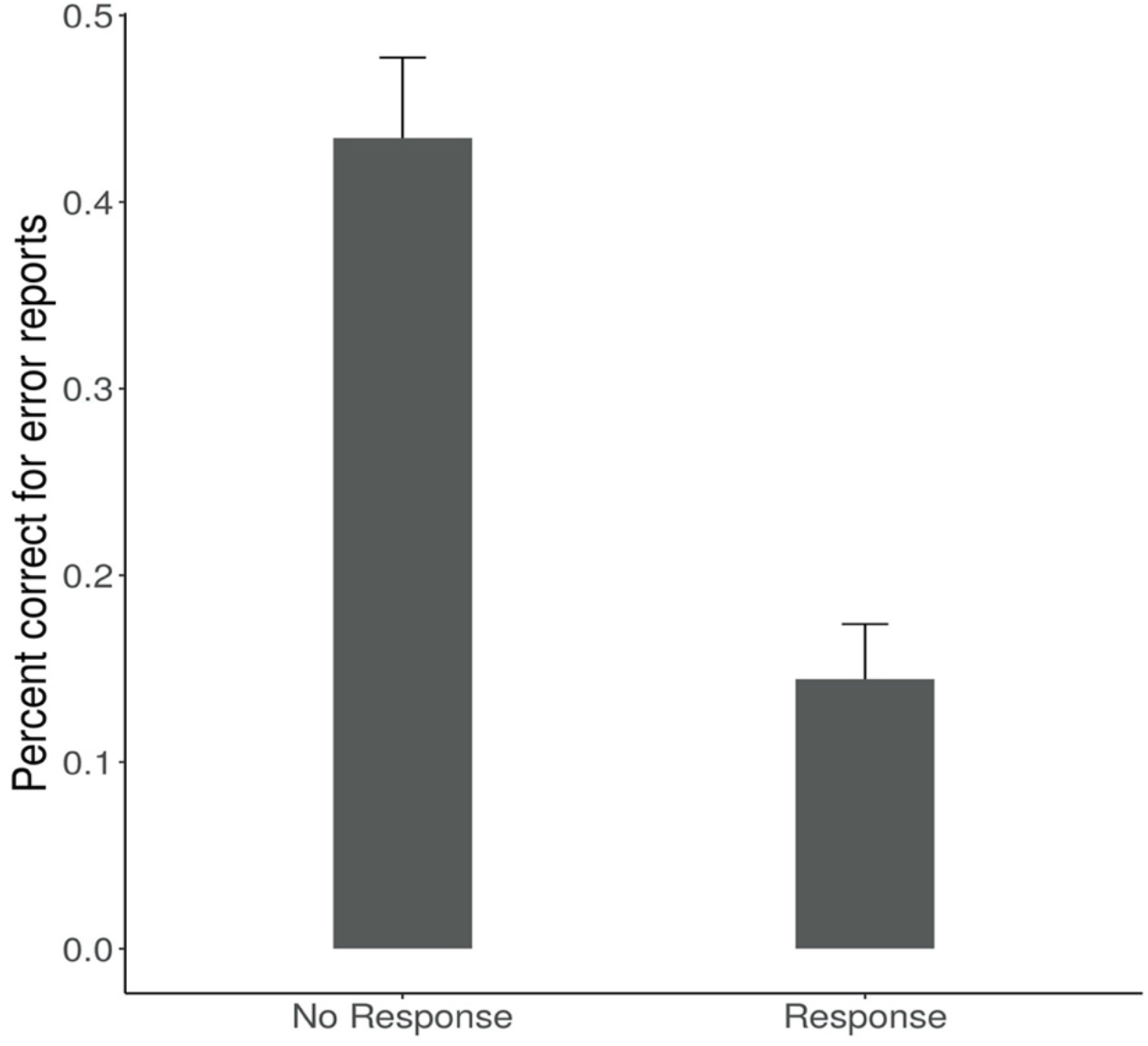
Percentage of correct responses when participants reported committing an error in each condition. The bars represent 95% confidence intervals.

### Confidence ratings

In the following analyses, we omitted trials in which participants reported errors (because a person could either rate their decision confidence or report an error). We compared levels of confidence between conditions using a linear mixed model with fixed and random effect of condition and participant-specific intercept. The analyses showed that reported confidence was higher in Response trials compared to trials in which participants did not respond (Table 2). When fixed and random effects of accuracy and its interaction with condition were also included in the model, we found that this effect was significant for both correct and erroneous responses (no detected interaction between accuracy and condition: *t*(52) = −0.13, *p* = .89).

**Table 2.**
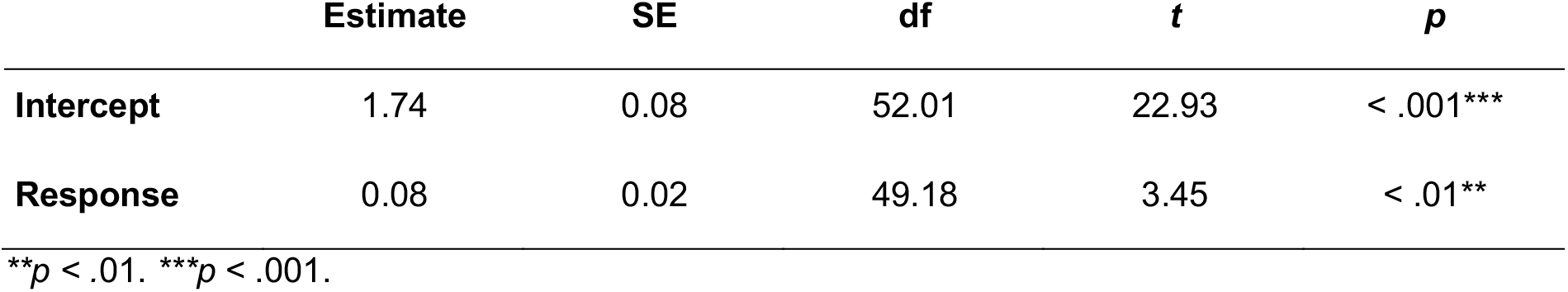
The linear mixed model estimation of the relationship between motor response and confidence level.

One potential confounder of the relation between confidence and response is task difficulty, which might influence both the probability of giving a response and the level of confidence. Therefore, we fitted another model in which we controlled for the dot ratio to find out whether the effect remains statistically significant. We assumed that a decision is easier when the difference between the number of dots on both sides is larger. We fitted a linear mixed model with confidence level as a dependent variable and fixed effects of condition, difficulty (3 levels), and all random effects. The model is presented in Table 3. Intercept refers to the No response condition and the lowest level of difficulty (easy). Response estimates the average effect of condition on confidence level independently of difficulty, and this relationship remains statistically significant. The other two coefficients estimate the difference in confidence between the lower level of difficulty and the two other levels; they show that confidence is significantly lower in medium and difficult trials compared to easy trials.

**Table 3.**
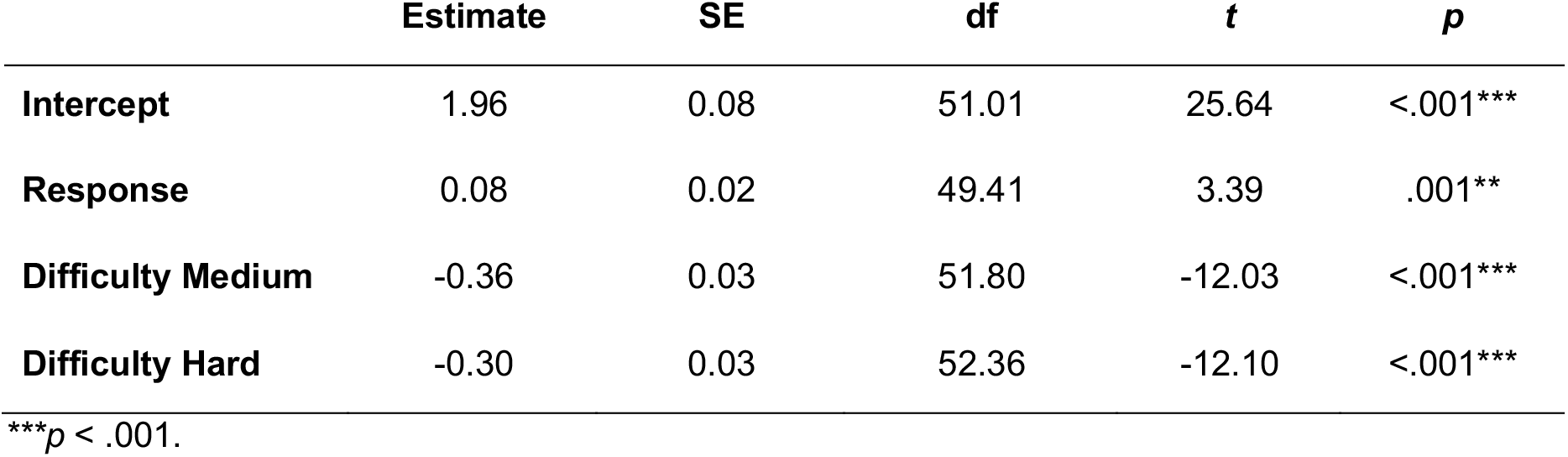
The linear mixed model estimation of the relationship between condition and confidence level controlled for difficulty level.

### The relation between decision accuracy and confidence judgments

In this analysis we tested whether the relationship between decision accuracy and confidence ratings differed between the two conditions by comparing the extent to which confidence ratings predicted accuracy in the decision task. Here we used logistic regression analysis because we did not want to rely on strong untested assumptions about the source of confidence that are implicit in popular alternatives based on signal detection theory (Paulewicz, Siedlecka, & Koculak, 2020).

We fitted a mixed logistic regression model to accuracy data with condition (2 levels), confidence (4 levels), and their interaction as fixed and random effects. To improve the readability of the regression coefficients, confidence ratings were centred on the lowest value (“guessing”), and the reference condition was the No response. Therefore, the regression slope reflects the relation between accuracy and confidence rating, while the intercept informs about the difference between chance level and decision accuracy when participants reported “guessing” in the No response condition.

The intercept was not significant, therefore we did not detect a difference between participants’ decision accuracy when reporting guessing and chance level in the No response condition (*z* = 1.92, *p* = .05). This lack of significant difference from chance level performance when reporting guessing was also not significantly related to condition (*z* = - 0.89, *p* = .38). However, the model revealed a statistically significant relationship between decision accuracy and confidence level in the No response condition (*z* = 13.64, *p* < .001), and this relationship was significantly stronger in the Response condition (*z* = 2.88, *p* < .01). The model coefficients are presented in Table 4, and model fit, together with percentage of correct responses for each confidence rating and frequency of each rating are presented in Figure 3.

**Table 4.** The mixed logistic regression model estimation of the relationship between accuracy and confidence level in both conditions.

|  | Estimate | SE | z | p |
| --- | --- | --- | --- | --- |
| **Intercept** | 0.12 | 0.06 | 1.92 | .05 |
| **Response** | -0.06 | 0.06 | - 0.89 | .38 |
| **Confidence** | 0.47 | 0.03 | 13.64 | < .001*** |
| **Confidence: Response** | 0.11 | 0.04 | 2.88 | < .01** |

\*\* $p < .01$ . \*\*\* $p < .001$ .

**Figure 3.**
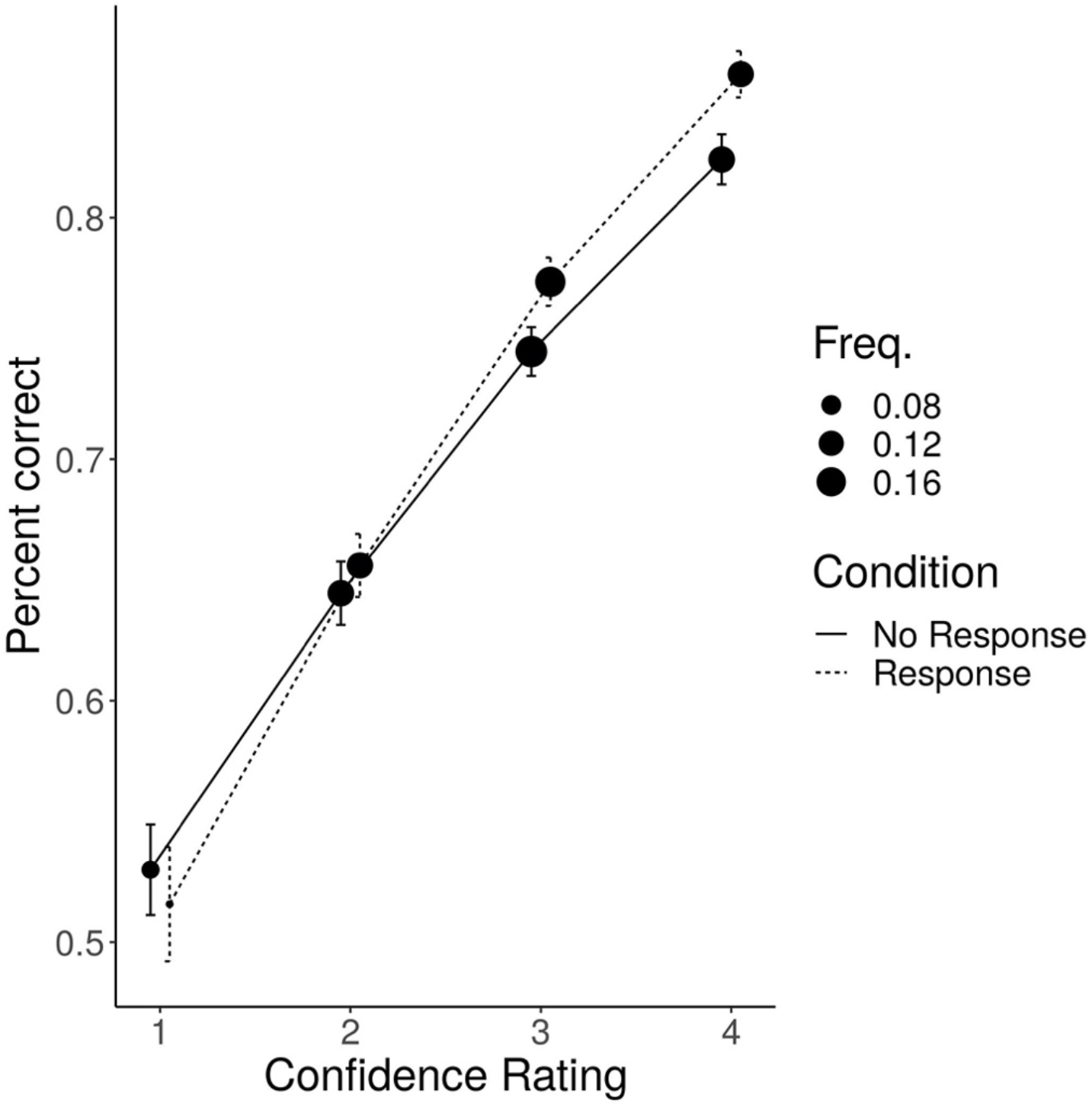
Model fit for the relationship between response accuracy and confidence ratings in each condition. The position of filled circles represents average accuracy for each scale point. The bars represent 95% confidence intervals. The frequency describes the proportion of each confidence rating.

### Response times

Finally, we analysed reaction times in the Response condition. Correct responses were given faster than incorrect responses (t(52.19) = −6.31, p < .001, a mixed linear model analysis with fixed and random effect of accuracy and participant-specific intercept). Slower responses were followed by lower confidence ratings (t(47) = −6.34, p < .001, a linear mixed model with fixed and random effect of confidence). Analysis of incorrect response times showed that reported errors were faster than unreported errors (t(5414) = 4.84, p < .001, a mixed linear model with random intercept).

## Discussion

In this experiment we compared confidence judgments and error reports between trials in which, depending on their decision about the stimulus, participants did or did not respond. The data showed that when given after a response, confidence ratings predicted decision accuracy better and reports of errors more often indicated an actual error. The data also showed that after responding participants reported higher confidence in both correct and incorrect decisions and independently of task difficulty. Additionally, analyses of reaction times showed that slower responses were followed by lower confidence ratings, while fast responses were associated with high confidence that a decision was correct or incorrect.

The results support previous findings showing that confidence judgments are more strongly related to decision accuracy when measured after a person responds (Pereira et al., 2018; Siedlecka et al., 2019; Wokke et al., 2019); this suggests that motor response provides metacognitive processes with additional information such as deliberation time, motor fluency, or internal accuracy feedback (Kiani, Corthell, & Shadlen, 2014; Susser & Mulligan, 2015; Ullsperger et al., 2010). Moreover, the results are consistent with findings from studies in which the occurrence of motor response was experimentally manipulated, showing that confidence in decisions is increased by action paired with decision-relevant stimulus characteristics (Filevich et al., 2019; Siedlecka et al., 2020). Since confidence was higher after correct and incorrect responses, this supports the interpretation that carrying out a response increases confidence by indicating successful completion of the decisional process (Siedlecka et al., 2020).

To our knowledge, this is the first study to compare the accuracy of error reports between decisions that were and were not followed by motor responses. The data showed that even though participants reported a similar number of errors in both conditions, their error reports were more accurate after they had responded. While over 80% of error reports indicated an actual error in Response trials, in No response trials participants were mistakenly convinced that they had made an error in almost half of the cases. As decision accuracy and confidence level were lower in trials without responses, this result suggests increased uncertainty: participants did not respond when they did not collect enough evidence for either choices in a given time, therefore they were also not able to accurately assess their performance. However, we would expect that in the case of high decision uncertainty, participants would report low confidence or guessing instead of an error (Scheffers & Coles, 2000). Studies on confidence, including this one, show that slower and more difficult decisions are associated with lower confidence (Desender, Van Opstal, & Van den Bussche, 2017; Kiani et al, 2014). At the same time, studies on error awareness indicate that errors are reported when sufficiently strong evidence (compared to noise and evidence for an already given response) passes the awareness threshold (Ullsperger et al., 2010; Wessel, 2012). Interestingly, even though there are theories about how and when error awareness emerges, not much is known about the possible causes of error illusions. On one hand, if a decision-related action increases confidence as an indicator that the decision process has completed, lack of action following the decision might lead to internal error signals. At the same time, without action, accuracy assessment is based on fewer sources of information. Moreover, in some cases this information might be unreliable: for example, when a chosen option is not associated with any action plan, even weak activation of a response linked to an unchosen option might bias the judgment towards an error report.

The results of this study are consistent with the view that both confidence judgments and error reports are informed by post-decisional processing. In most confidence models, however, post-decisional processing refers to ongoing decision-related evidence and not to action-related information (e.g. Moran, Teodorescu, & Usheret, 2015; Pleskac & Busemeyer, 2010). One explanation of the role of action in metacognitive judgments is offered by the hierarchical Bayesian model of self-evaluation, according to which such judgments are based on second-order inference about the performance of the decision system. Crucially, decision and confidence are computed separately in this model, and action provides information about the decision process that is not otherwise accessible to the metacognitive system (Fleming & Daw, 2017). We propose that confidence and error judgments are the results of metacognitive processes that consist of multiple monitoring and regulating loops that occur at consecutive stages of stimuli identification, decision making, and action preparation and execution (Paulewicz, Siedlecka, & Koculak, 2020). Therefore, each stage of this process, before and after action, is monitored and might trigger regulation. For example, studies have shown that the monitoring system can detect, inhibit, and correct erroneous response activation (so-called “partial errors”) before the response is given, which therefore helps avoid error commission (Burle, Possamaï, Vidal, & Bonnet, 2002; Scheffers, Coles, Bernstein, Gehring, & Donchin, 1996).

The limitation of this study is its observational assessment of the relation between confidence level and action. Although confidence is often assumed to reflect the assessment of the whole decisional process, it might determine the outcome of this process, and more importantly, if and how people will act on their decisions. The choice of action might be affected by the results of monitoring the decision process. In our study, participants might have chosen to respond when they felt confident in their decision and did not respond when their uncertainty was high. Therefore, the confidence report at the end of a trial might have been affected by the response and by the level of confidence preceding that response. However, confidence that one has committed an error can only arise after that decision or action has been made. It is still plausible that the differences in the accuracy of error reports were due to different types of errors, which were in turn related to pre-decisional confidence. While mistakes in No response trials could have mostly resulted from high uncertainty about stimulus characteristics, in Response trials we might have observed slips and premature responses. However, as in our procedure there was no prepotent response, this explanation suggests that incorrect responses were first given quickly because of high confidence in decisions, and then this confidence changed into confidence that an error had been committed. It would be informative in future studies to find a way of probing subjective confidence in different stages of the decision-making process to find out how it changes before and after decisions and actions, or how it predicts commitment to action.

To summarize, the results of this study support the view that action plays an important role in shaping subjective confidence and allows better assessment of decision accuracy.

## Author contribution

MS & MK conceived the plan of the study. MK prepared the experimental procedure and ran the tests. BP analysed the data in collaboration with MS. MS wrote the manuscript, all authors provided comments.

We would like to thank Justyna Hobot and Magdalena Senderecka for commenting on the manuscript, and Kinga Ciupińska and Alicja Krzyżewska for their help with data collection.

